# Decoding the gene co-expression network underlying the ability of *Gevuina avellana* Mol. to live in diverse light conditions

**DOI:** 10.1101/240572

**Authors:** E Ostria-Gallardo, A Ranjan, Y Ichihashi, LJ Corcuera, NR Sinha

## Abstract

- *Gevuina avellana* Mol. (Proteaceae) is a typical tree from the South American temperate rainforest. Although this species mostly regenerates in shaded understories, it exhibits an exceptional ecological breadth, being able to live under a wide range of light conditions. Here we studied the genetic basis regulating physiological acclimation of the photosynthetic responses of *G. avellana* under contrasting light conditions.
- We analyzed carbon assimilation and light energy used for photochemical process in plants acclimated to contrasting light conditions. Also, we used a transcriptional profile of leaf primordia from *G. avellana* saplings growing under different light environments to identify the gene co-expression network underpinning photosynthetic performance and light-related processes.
- The photosynthetic parameters revealed optimal performance regardless of light conditions. Strikingly, the mechanism involved in dissipation of excess light energy showed no significant differences between high and low-light acclimated plants. The gene co-expression network defined a community structure consistent with the photochemical responses, including genes involved mainly in assembly and functioning of photosystems, photoprotection, and retrograde signaling.
- Our ecophysiological genomics approach provides an understanding of the molecular regulatory mechanisms that allows this tree to have an optimal balance between photochemical, photoprotective and antioxidant performance in the diverse light habitats it encounters in nature.

## Introduction

Effective collection of light is a vital task for photosynthetic organisms. It is achieved by the development of several strategies to maximize the efficiency of this process (Janik et. al., 2015). Under low light conditions, plants need to harvest most available photons to sustain photosynthesis, while under high light they must dissipate the excess energy absorbed in order to avoid photodamage (Ruban 2016). Thus, in ecosystems where light is the most dynamic and limiting resource, its availability determines the establishment, growth, and survival of plant species (Valladares et. al., 2002). For example, in evergreen forests light availability is highly variable due to a number of factors such the rotation of the earth, wind-induced movement of leaves, leaf deployment, canopy architecture, and dynamics of gaps formation (Whitmore 1989; Valladares et. al., 1997). Also, plants, and particularly trees, are progressively exposed to higher irradiances as they grow taller (Kira and Yoda 1989; Coopman et al. 2011). Consequently, several tree species can, to some degree, change their light requirements through a suite of anatomical, morphological, and physiological leaf traits that allow them to adjust the photosynthetic apparatus to suit a new light environment (Coopman et al. 2008; Lusk et al. 2008; Poorter et al. 2010; Coopman et al. 2011).

Plant adaptation to a specific light environment can be viewed in terms of benefits (e.g., photosynthetic carbon gain) and cost (e.g., maintaining morphological and physiological flexibility) of various traits. Adaptation is expected to result in a situation where the ratio of benefits to cost is maximized (Björkman 1981). In shade environments, there is little return on investment in increasing the capacity of photosynthetic reactions, and resources are better invested in light harvesting. The reverse is true in high light environments, where electron transport, carboxylation capacity, and stomatal conductance tend to be maximized. After Givnish (1988) presented a comprehensive view of plant adaptations to sun and shade, a vast number of studies have described several morpho-anatomical and biochemical traits with adaptive value to optimizing the use of light. For example, the light harvesting complexes (LHCs) respond to the amount of light by having a dual functionality based on the different configurations of the pigment-protein system of the complex, resulting in a switch from light-harvesting to a photoprotective state (Liguori et. al., 2015). Nevertheless, there is still little information about the genetic aspects governing the photochemical and photoprotective responses of plants to fluctuating light.

Canopy shade changes light quality and quantity available to understory plants. Thus, inside the canopy a mosaic of light environments exists that strongly affects plant growth and fitness. Light perception and subsequent transduction events can converge on shared molecular pathways to elicit a proper response, specially in new forming leaves, to optimize the photosynthetic process and competitiveness between individuals (Franklin 2009; Smékalová et al., 2014). Key aspects of the synchronicity of growth responses to light environment are: (i) the integration of multiple external and internal cues; (ii) the uses of shared regulatory elements such as transcription factors, target proteins, hormones and secondary metabolites in diverse developmental cascades; and (iii) the role of physiological feedback to provide environmental information to the new forming photosynthetic tissues (Sultan 2010). This ultimately constitutes a critical aspect of the ecological breadth for forest species. For these reasons, plants have developed life strategies from which it is possible to recognize two major groups, shade-tolerant species and light-demanding species. The morphological and biochemical traits of the two types of species have been optimized for the light range within their ecological breadths.

An exceptional case of broad ecological breadth is seen in *Gevuina avellana* Mol. This early successional and short-lived temperate rainforest tree species is endemic to Chile and Argentina. Although this species mostly regenerates in shaded understories (Donoso 1978), seedlings and saplings can be found growing under a wide range of light conditions, from less than 5% of canopy openness to ca. 50% (Lusk 2002 and references therein) as well as in forest edges (personal observation, Fig. 1). Its leaves are highly plastic (Donoso 2006). Among the ca. 13 coexisting tree species in this forest-type (Lusk et al. 2008), *G. avellana* is the only known species that shows strong heteroblasty. Its leaves change drastically from simple to highly compound leaves during early development (Ostria-Gallardo et. al., 2015). In addition, when growing under high light conditions, this species produces larger and more complex leaves (Fig. 1). We previously demonstrated that the heteroblastic trajectory of leaf size and complexity depends on the prevailing light availability of the specific micro-environment where individual plants are established and grow (Ostria-Gallardo et al., 2015, 2016). This involves the coordination of light-mediated signals, hormone synthesis and signaling, and the heterochrony of the ontogenetic program. The responses of functional traits, such as leaf longevity and leaf mass per area, to light availability and the ontogenetic variation in light requirements have been reported for this species (Lusk et al., 2008; 2011). However, the genetic basis that regulates the physiological acclimation allowing *G. avellana* to grow in a wide variety of light environments remains unknown. Our hypothesis is that the ability of *G. avellana* to grow under a wide range of light conditions relies on the flexibility in the gene network to respond appropriately and comprehensively to signals from the prevailing light environment. We used two experimental approaches to identify gene co-expression networks that underpin the photosynthetic and metabolic performance allowing this tree to develop under different light environments. We used the transcriptome dataset of leaf primordia from *G. avellana* previously published by Ostria-Gallardo and coworkers (2016) to conduct a gene co-expression network analysis with genes involved in photosynthesis and light-related processes. In parallel, we conducted a growth chamber experiment in which saplings of *G. avellana* were acclimated to high and low light conditions in order to evaluate the effect of light intensity on the responses of gas exchange and light energy use.

**Fig. 1.**
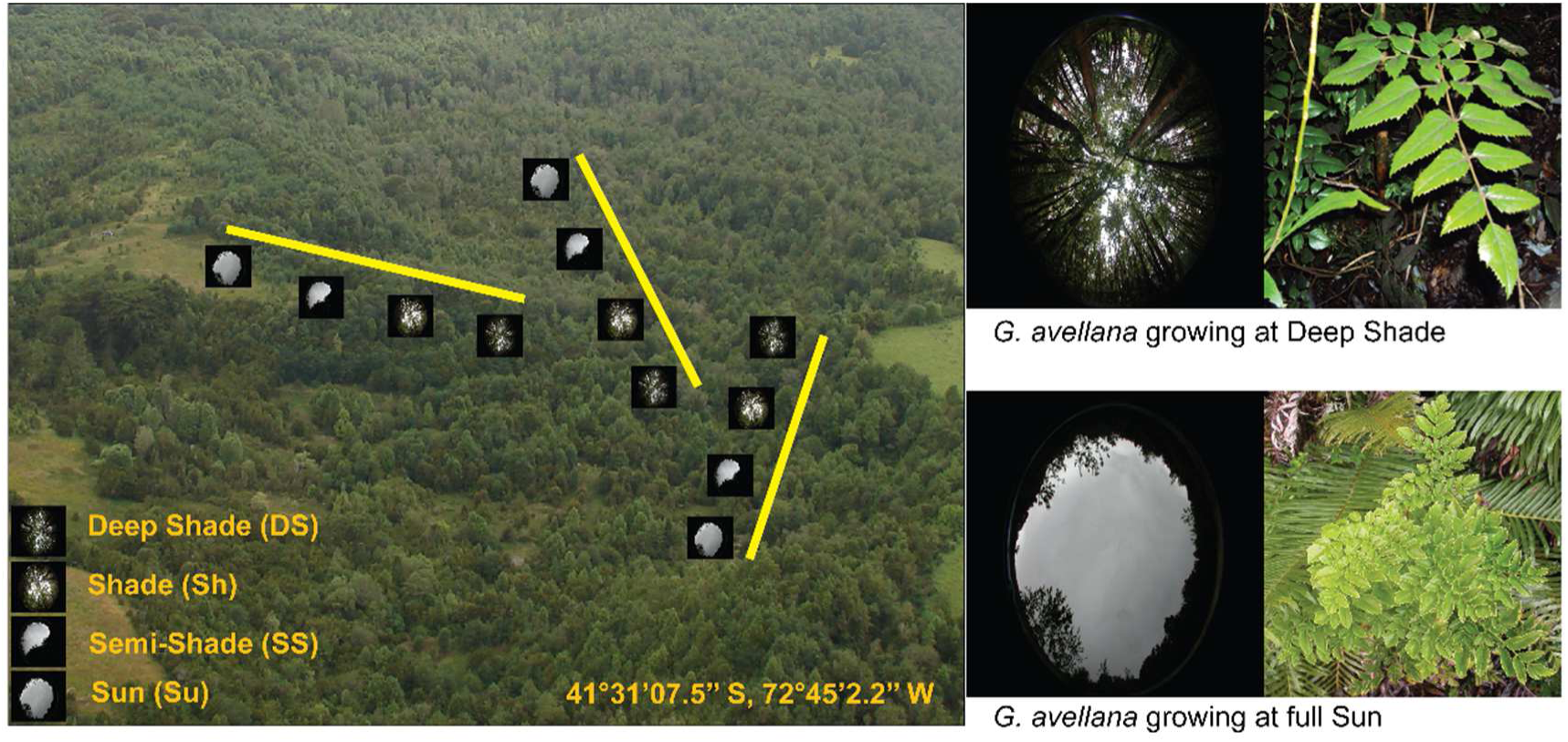
At the left, an aerial view of the study site showing the three areas of leaf primordia from *G. avellana* sampling for saplings growing in different light environments. At the right, the phenotype of *G. avellana* leaves growing under deep shade (DS) and full sun conditions (SU). (Modified from Ostria-Gallardo *et. al*., 2016).

## Materials and Methods

### Field and controlled-environment experiments

Field experiment was conducted in a secondary temperate rainforest stand located in South Central Chile (Katalapi Park: 41°31′07.5″ S, 72°45′2.2″ W; for further climatic details and forest structure see Coopman *et al*., 2010 and Ostria-Gallardo *et al*., 2015). Sampling and light environment characterization was made as described in detail in Ostria-Gallardo et al. (2016). Briefly, three randomly selected areas of 100 m long and 4 m wide cover most of the light gradient across the forest canopy (Fig. 1). The light environments along the selected areas were analyzed by a rigorous quantitative method to determine the percentage of canopy openness (%CO) by using the Gap Light Analyzer 2.0 software (GLA, Frazer *et al*., 1999). Thus, four light environments were established by using interquartile ranges of the %CO (deep-shade = DS; shade = Sh; semi-shade = SS; and sun = Su). Within each light environment, the healthiest seed-grown *G. avellana* plants were carefully selected ranging from 4 to 200 cm in height and the total chlorophyll content was measured with a chlorophyll meter SPAD-502 (Konica Minolta Corp. Sensing Europe B.V). Given height differences of plants within each light environment, we characterized the light availability above each individual and then the leaf primordia from the apex were collected in liquid nitrogen and stored at -80°C.

In the controlled-environment experiment, we used one-year old *G. avellana* plants grown from seed, and of the same height. Plants were carefully transplanted to black plastic bags filled with organic soil, irrigated at soil field capacity and fertilized twice with a 0,4 gl^-1^ of a commercial solution (Phostrogen Plant Food, NPK ratio of 10:14:27 plus micronutrients). Plants were acclimated in nursery conditions for about 3 months, in which, at sunny day, the minimum and maximum photosynthetic active radiation (PAR) was ca. 3 and 400 µmol m^-2^s^-1^, respectively. To study light acclimation of new expanding leaf, after nursery acclimation, a total of 14 pots were placed in a controlled-environment grow chamber at 15°C in a 15-h photoperiod for three months under two light treatments; 7 pots under low light (ca. 100 µmol m^−2^s^−1^) and 7 pots under high light (ca. 1000 µmol m^−2^s^−1^). The light sources were metal halide lamps coupled with a cooling system to maintain leaf temperature around 15 to 20 °C.

### Gas exchange and use of light energy

Light curves ranging from 0 to 2000 µmol m^−2^s^−1^ were measured in attached leaves to determine the maximum CO_2_ assimilation by using a Portable Photosynthesis System (LI-6400; LI-COR Inc., Lincoln NE, USA). CO_2_ reference concentration was 400 ppm with a flux rate of 350 ml min^−1^. Leaf temperature inside the chamber was maintained at 20 °C and 65–75 % of relative humidity. Dark respiration rate (*R*_d_), photosynthetic capacity (*A*_max_), instantaneous water use efficiency (*WUEi*) and light compensation and saturation points were determined for plants acclimated to high and low light. All measurements were performed between 0900 and 1400 h. Gas exchange values were adjusted for the cuvette area/actual leaf area ratio. In the same set of attached leaves, we measured the fluorescence signals of chlorophyll *a* by a pulse-amplitude modulated fluorimeter (FMS2, Hansatech Instruments, King’s Lyn, U.K). Leaves were dark acclimated for 30 min. at 15°C. Different light pulses were applied during 3 min. following standard settings of the fluorometer (see detailed information of light pulses in Coopman et al., 2008). We calculated electron transport rate [ETR(II)], quantum yield of PSII[ϕ (II)], and yield of energy dissipation by antenna down-regulation [ϕ(NPQ)] according to Kramer *et al* (2004). Data was analyzed after check normality assumptions under an ANOVA test with P value ≤ 0.05. When data did not meet the normality assumptions, we used the non-parametric test of Kruskal-Wallis.

### Principal Component Analysis (PCA) and Self-Organizing Maps (SOM) Clustering

In the present study, we used the transcriptional profile of *G. avellana’s* leaf primordia under different light environments reported in Ostria-Gallardo et al., 2016 (refer this citation for details of isolation of mRNA, pipeline for library preparation, assembly and annotation of transcripts). Normalized RSEM-estimated counts were used for clustering assembled ORFs based on expression patterns (Chitwood *et al*., 2013). In order to detect the effects of light availability on gene expression, we selected genes from the upper 75% quartile of coefficient of variation for expression across light environments. The scaled expression values within samples were used to cluster these genes for a multidimensional 2 × 3 hexagonal SOM using the Kohonen package on R (Wehrens & Buydens, 2007). 100 training interactions were used during clustering with a decrease in the alpha learning rate from ca. 0.0050 to 0.0035 (Supplemental Fig. S1). SOM outcome was visualized in PCA space where PC values were calculated based on the gene expression of samples across light environments. This first clustering process was a necessary step to select groups of genes highly correlated to light environment and involved in photosynthesis and light-related processes.

### Gene Co-expression Network Analysis

From the PCA-SOM clustering method, we selected a total of 436 subset of annotated genes functionally relevant in photosynthesis and light-related responses/processes (Supplemental Dataset 1) to perform a weighted gene coexpression network analysis by using the Weighted Gene Coexpression Analysis (WGCNA; Langfelder and Horvath 2008) and the igraph R packages (Csardi and Nepusz 2006). In order to meet the scale-free topology criteria for WGCNA, we use a soft threshold (β) value of 17 for the calculation of the adjacency matrix. Network properties such as connectivity, centralization, modularity and community structure was calculated for each of the using the resources and algorithms of the fastgreedy.community and the igraph R package. Finally, we used customs graphs functions of the igraph packaged for network visualization.

## Results

### Light acclimation effects on maximum photosynthetic capacity and light energy partitioning

The light response curves for gas exchanges in *G. avellana* saplings showed no significant differences between low-light (LL) and high-light (HL) acclimated plants (*P* = 0.15). However, when considering only light intensity ≥ 500 µmol m^−2^s^−1^ the values of A_max_ of the sun acclimated saplings were significantly higher (*P* = 0.004) (Figure 2). The light compensation point was the same in shade and sun acclimated plants but in the latter, the light saturation point was twice as high. Evapotranspiration per unit area was similar between both groups of plants (Figure 3a). Despite the similar rates of water loss, WUE_i_ in sun acclimated plants was significantly higher (*P* = 0.05) in comparison to those acclimated to shade (Figure 3b).

**Fig. 2.**
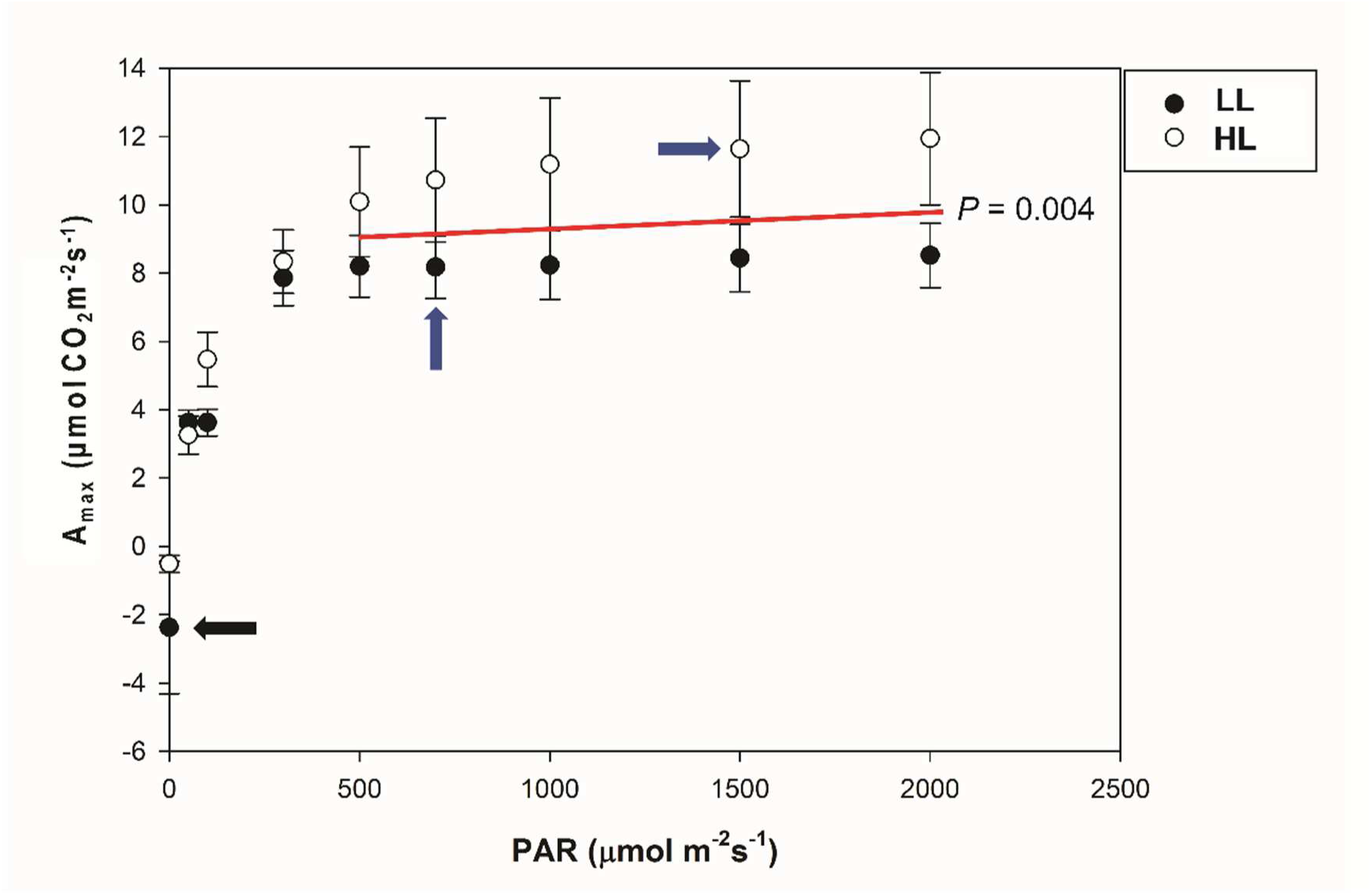
Light response curve of the photosynthetic capacity (A_max_) in saplings of *G. avellana* acclimated to low-light (LL, closed circles) and high-light (HL, open circles). Continuous red line denotes the part of the curve in which the difference in A_max_ of LL and HL plants turns significant at *P* < 0.05 (from 500 to 2000 pmol m^−2^s^−1^). Both species show the same light compensation point (black arrow), but differ in their light saturation point (blue arrows).

**Fig. 3.**
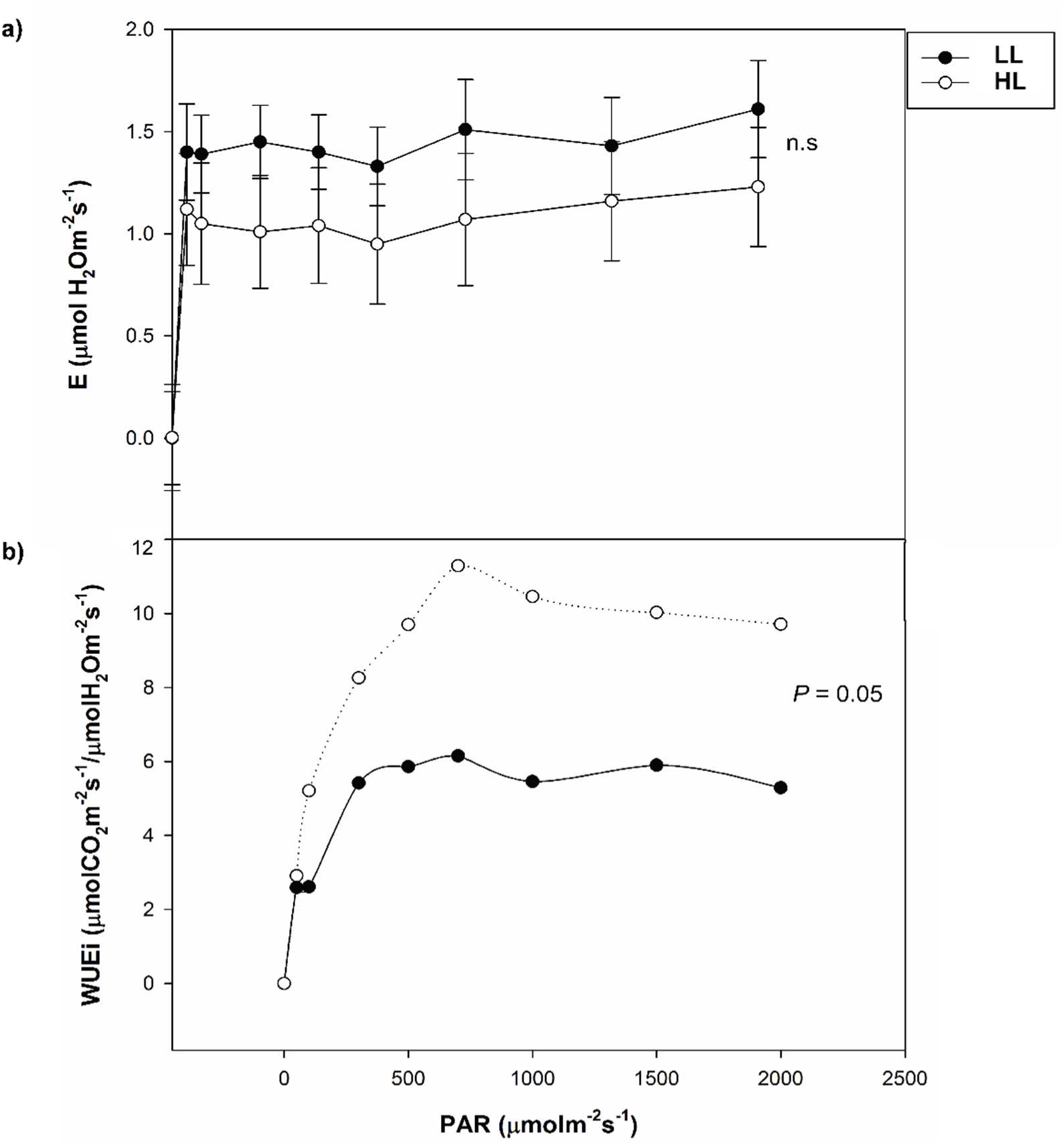
(a) Evapotranspiration (E) and (b) instantaneous Water Use Efficiency (WUEi) curves in response to increasing light. (a) shows the light response curves of E for low-light (LL) and high-light (HL) acclimated plants without evidence of statistical differences (n.s), indicating a similar pattern of water loss. (b) shows the light response of WUEi for LL and HL acclimated plants showing significant differences between both treatment (*P* = 0.05), indicating that HL acclimated plants are able to optimize the ratio of Carbon intake to water loss.

There was no difference in the total relative chlorophyll content in leaves of shade and sun acclimated saplings (Figure 4a). Fluorescence of chlorophyll *a* revealed that, despite differences between low and high light acclimated plants, all performed well in terms of photochemistry. The maximum quantum efficiency of photosystem II (Fv/Fm, Figure 4b) was higher in low light acclimated plants (mean=0.87, *P* = 0.0357). However, the actual quantum yield of photosystem II (ΦPSII), the electron transport rate (ETR) and the photochemical quenching (qP) increased significantly in plants growing at high light *(P* < 0.05; Fig. 4 c, d and e). Contrary to our suppositions, the values for non-photochemical quenching (NPQ) showed no significant differences between plants from the two light conditions (P = 0.39; Fig. 4f).

**Fig. 4.**
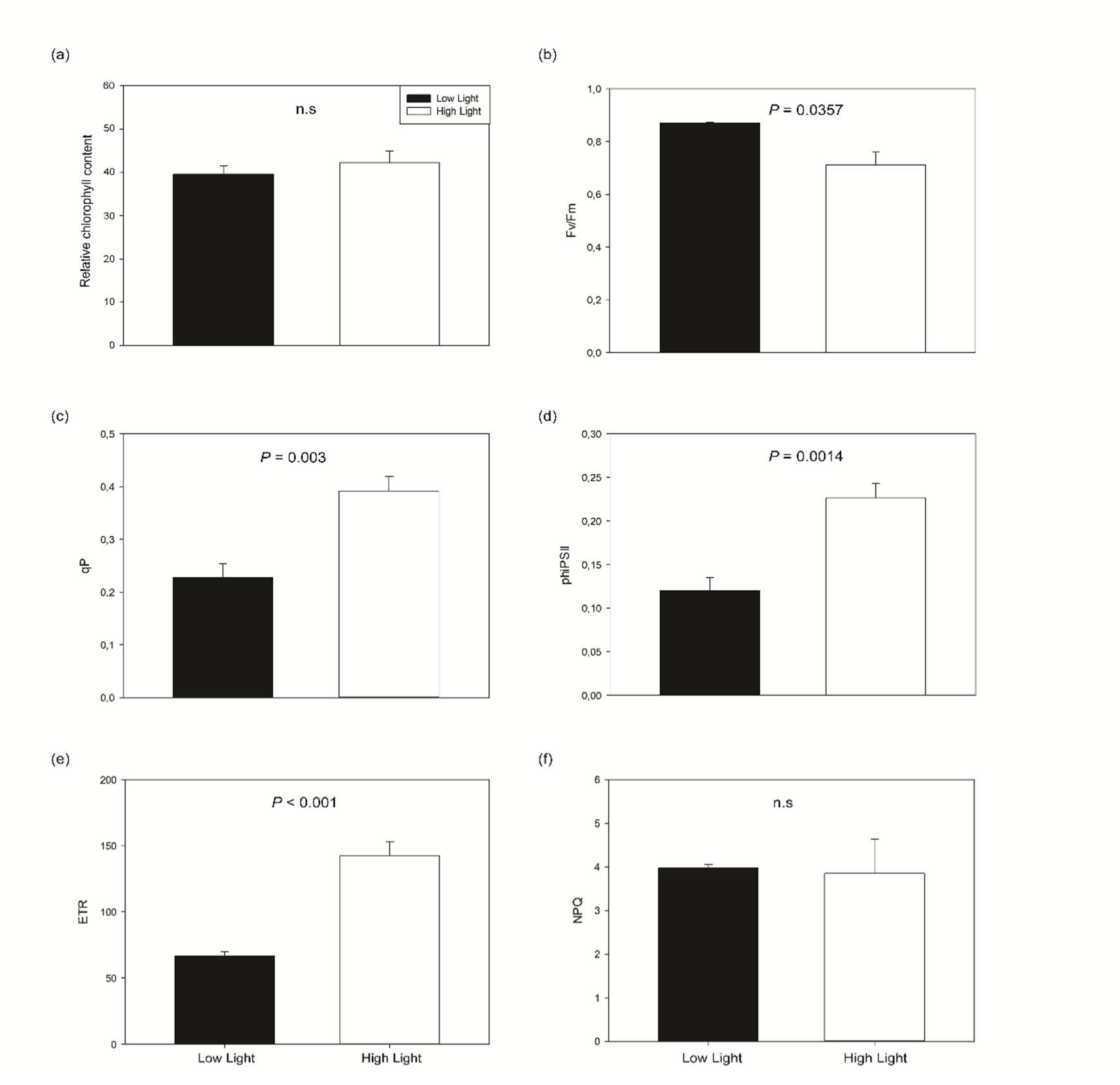
Relative chlorophyll content and parameters of fluorescence of chlorophyll *a*. (a) The relative chlorophyll content showed no significant differences (*P* < 0.05) between low-light and high-light saplings (total n = 40). Among fluorescence parameters, the maximum quantum efficiency (Fv/Fm, (b)) showed significant differences between saplings acclimated to low and high-light, showing higher values in those that are low-light acclimated. The photochemical quenching (qP, (c)), the actual quantum yield (phiPSII, (d)), and the electron transport rate (ETR, (e)) showed significant differences between saplings acclimated to low and high-light, with all these parameters being higher at high-light. The non-photochemical quenching (NPQ, (f)), involved in photoprotective processes, showed no significant differences between saplings. For the fluorescence of chlorophyll *a*, all statistical significances were at *P* < 0.05 with n=11 and n = 7 for low and high-light acclimated saplings, respectively.

### *Gene coexpression network of* G. avellana *in response to light environment*

In order to analyze gene expression changes induced by light quality, we subsampled data from the available transcriptome for *G. avellana* (Ostria-Gallardo *et. al*., 2016). A Principal Component Analysis conducted on the full dataset of differentially expressed genes (DEG), coupled with a Self-Organized Map clustering (PCA-SOM), revealed patterns of transcript accumulation, and variation across the light gradient (i.e., deep shade = DS; shade = Sh; semi shade = SS; sun = Su; see Material and Methods for details) explained 75% seen in the data (Fig.5). Among these patterns, a clear separation between the shadiest (DS, cluster 2) and the sunniest (Su, cluster 6) light environments was observed in the PCA-SOM. We determined GO enrichment to infer biological relevance in each cluster. Genes in cluster 2 associated with DS were highly enriched in the functional categories of photoperiod-related processes. Transcripts in clusters 1 and 3, associated with Sh and SS, respectively, were enriched for the functional categories of cell proliferation, gene regulation, signaling and development. Transcripts in cluster 6 were enriched in translational machinery. From these SOM clusters (1, 2, 3, and 6), we carefully reviewed the annotated genes (see Supplemental Dataset 1) with high scaled expression associated to each light environment, and then we selected a subset of 436 annotated genes engaged in photosynthesis and light-related responses/processes to construct a weighted coexpression network (see Materials and Methods for details). The resultant network was fitted to a scale-free topology criterion reasonable for biological networks (R^2^ = 0.72, slope=-0.72; Fig. 6 a and b). Based on the Fast Greedy modularity optimization algorithm, the network had 86 modules, a total of 11,752 edges and a modularity of 0.43. The network density (i.e., the portion of potential connections that are actually connected) was 0.12, whereas the mean of the shortest distance between each pair of genes in the network was 1.85.

**Fig. 5.**
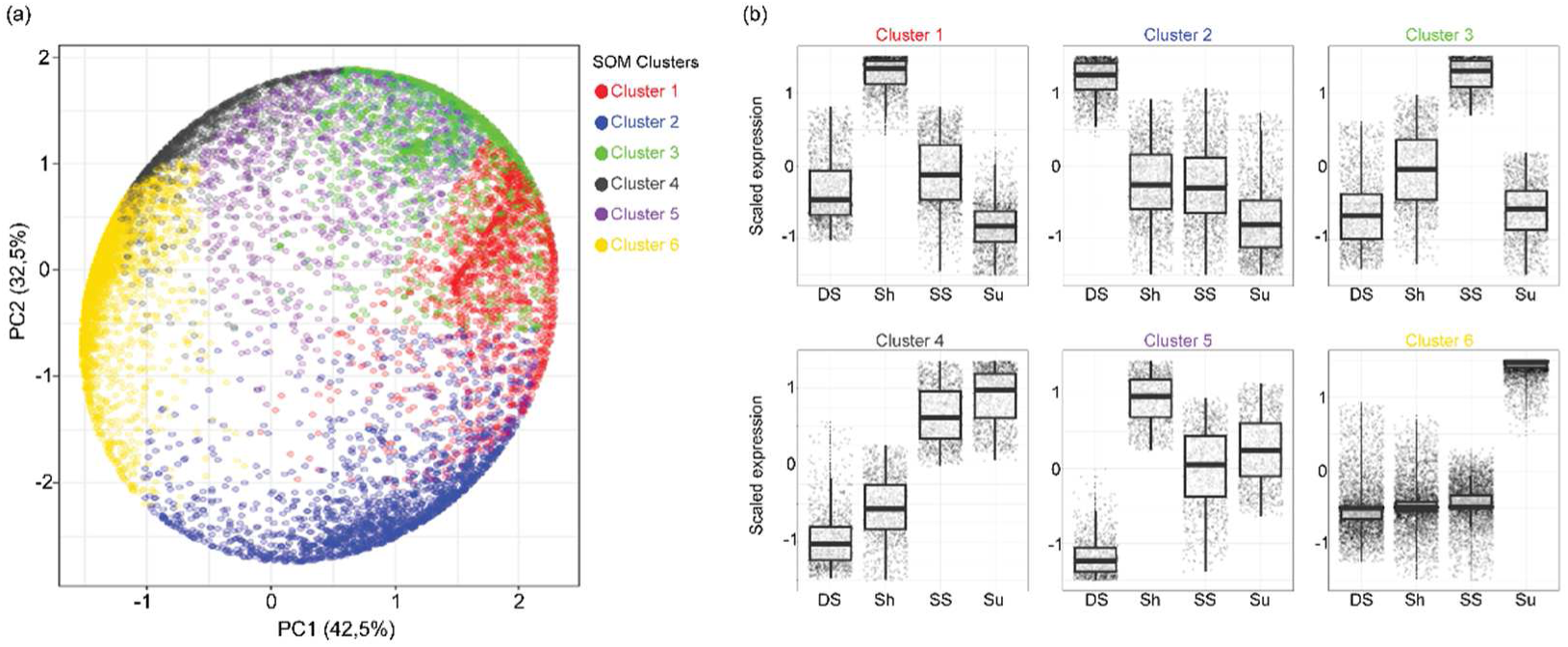
(a) Principal component analysis (PCA) with self-organizing map (SOM) clustering for gene expression in response to light environment classes. The PCA-SOM space shows two principal component (PC1 and PC2) that explain the higher percentage of variance in the dataset, and represents the clustering patterns of the transcriptional profile in response to light (indicated by different colors) (b) Box-plots show in detail the specific accumulation pattern of transcripts foe each light environment class within each cluster defined by the PCA-SOM. In each box-plot the horizontal line indicates the median while bars represent the maximum and minimum values of the scaled transcript abundance. Transcripts showing the highest accumulation pattern in one of the light environment classes from clusters 1, 2, 3, and 6 were used to select those transcripts highly involved in photosynthesis and light-related processes for further co-expression network analysis.

**Fig. 6.**
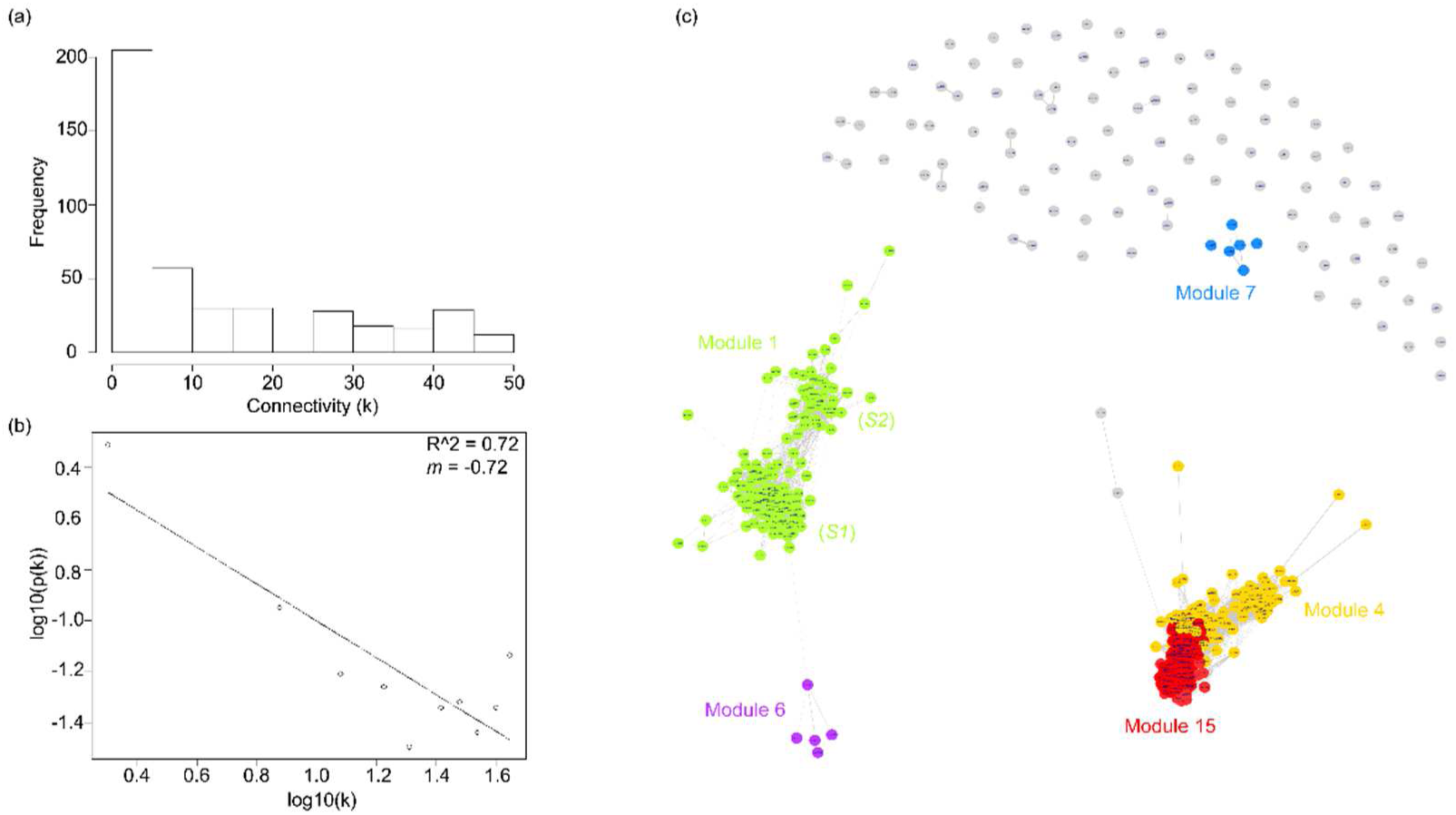
(a) The frequency distribution of connectivity. The histogram shows a large number of low connected genes and a small number of highly connected genes. (b) The log-log plot based on connectivity shows that the predicted co-expression network follows a scale free topology criterion (R^2^ = 0.72, slope = -0.72). (c) Gene co-expression network using transcript from clusters 1, 2, 3, and 6 (see Fig. 5b). The network highlights five of a total of eighty-six modules showing the denser connections between nodes. Each module (indicated by different colors) include transcripts with specific function in photosynthesis and light related processes. Module 1 contains transcripts encoding for Light Harvesting Complexes (LHC) and Photosystems proteins (*S1*), as well as enzymes involved in redox reactions of the Calvin cycle and reactive oxygen species (ROS) scavenging (*S2*). Module 4 contains transcripts involved mainly in the biosynthesis of carotenoids and terpenoids. Module 15 contains transcripts encoding enzymes involved in ROS scavenging by the ascorbate-glutathione cycle, and enzymes involved in glucose catabolism. Module 6 contains transcripts encoding proteins involved in photoprotection. Module 7 contains transcripts involved in terpenoids and terpenes biosynthesis and metabolism. Further details in the Result section and in Supplemental dataset 1.

Given the community structure of the network (composed of 86 modules), we focused on those modules showing a module membership *(MM*, i.e., genes highly connected with a module) greater than 3, resulting in 5 modules of interest. The highest *MM* was found in modules 1 and 4, with 125 and 124 genes respectively (Figure 6c). Module 15 has a *MM* of 79 genes, whereas module 6 and 7 had a *MM* of 5 and 6 genes respectively. An overview of module 1 reveals a coexpression of genes involved in photosynthesis (light reactions and carbon fixation), carotenoid and flavonoid biosynthesis. In specific, the community architecture of module 1 presents two well defined subgroups, *S1* and *S2* (Figure 6c). *S1* contained genes related to the assembly and photoprotection of the antenna complexes and photosystems (e.g., *PHYTOENE SYNTHASE, HIGH CHLOROPHYLL FLUORESCENCE PHENOTYPE 173, PHOTOSYSTEM II SUBUNIT O-2, LIGHT-HARVESTING CHLOROPHYLL-PROTEIN COMPLEX 1 SUBUNIT A4)*, whereas *S2* contained genes related to carbon reactions and also retrograde signaling (e.g., *GLYCERALDEHYDE-3-PHOSPHATE DEHYDROGENASE OF PLASTID 1, TRANSKETOLASE, RIBULOSE BIPHOSPHATE CARBOXILASE OXYGENASE SMALL UNIT, AAA-TYPE ATPASE)* (Supplemental Dataset 1, Supplemental Fig. S2a). An intermodular connection between module 1 and 6 reveals similar functional mechanisms of these modules, because the clustered genes within module 6 (*MM* = 5) are involved in photoprotection and glucose signaling (e.g., *EARLY LIGHT INDUCED PROTEIN 1, PROTON GRADIENT REGULATION 5, HEXOKINASE LIKE* 1). In addition, according to the light environment, we observed that genes involved in processes such the stabilization of trimeric forms of the antenna complex (e.g., CHLOROPHYLL *a-b* BINDING PROTEIN OF LHCII TYPE 1-LIKE) and minor antenna complexes *(CHLOROPHYLL a-b* BINDING PROTEIN CP26, *CP29)* are downregulated when plants are under full sun conditions (Supplemental Fig. S2b), suggesting regulation of antenna size and photoprotective function.

Next, within module 4 we found genes related to the oxidative damage regulation (i.e., *THIOREDOXIN-DEPENDENT PEROXIDASE)*, and especially in the glutathione-ascorbate cycle (e.g., *CATALASE1. ASCORBATE PEROXIDASE 3, MONODEHYDROASCORBATE REDUCTASE, GLUTATHIONE REDUCTASE)* (Supplemental Fig. S2c). This module was tightly connected with module 15 (*MM* = 79) whose components are engaged in metabolic pathways such as the control of chlorophyll content, photorespiration, lipid peroxidation and respiration (e.g., *PROTOCHLOROPHYLLIDE OXIDOREDUCTASE A, GLUTAMATE:GLYOXYLATE AMINOTRANSFERASE 1, GLYOXYLATE REDUCTASE, ATB2, ATPase, ISOCITRATE DEHYDROGENASE V, SUCCINATE DEHYDROGENASE 2–2*) (Supplemental Fig. S2d). Finally, genes contained in module 7 (MM = 6) function in the formation of parent terpene carbon skeletons (e.g., *TERPENE SYNTHESE 3, TERPENOID CYCLASE)* (Supplemental Dataset 1), and make up an isolated community without connection with other modules.

## Discussion

Using gas exchange and chlorophyll *a* fluorescence parameters we showed that in spite of been considered a semi-shade species, the basal eudicot tree *Gevuina avellana* has the physiological capacity to inhabit a wide range of light environments, from deep shade to sun, without significant effect on light energy use for photosynthesis. According to reports of Lusk and coworkers (2006, 2008) detailing the ecological effects of light availability on the light requirement of temperate rainforest tree species, those species that possess a longer leaf life span and that tend to accumulate multiple foliage cohorts are more stable to variations in light availability. *G. avellana*, particularly, does not have significant changes in minimum light requirement along its ontogeny (plants of 10 to 120 cm height; Lusk et al., 2008). This means that while growing, this tree may tolerate low light conditions without facing carbon starvation, as is seen to occur in shade intolerant species (Escandón et al. 2013). According to our results, the maximum quantum yield of low and high light acclimated plants reflects healthy photosynthetic apparatus capable of using most of the light energy to conduct effective photochemistry. Surprisingly, non-photochemical quenching (NPQ), which is a molecular response of photosynthetic membranes in which the excess of light is dissipated as heat (Ruban, 2016), showed similar values between low and high light acclimated plants. Since along the forest canopy in which *G. avellana* grows, different scenarios of high irradiance occur (i.e. high frequency of sunflecks, gaps or high light intensity in open space), this protective mechanism against photodamage is used by *G. avellana* in a balanced way, no matter the light condition, so that the use of light available for photosynthesis is not affected. We observed that below a photosynthetic photon flux density (PPFD) of 500 µmol m^−2^s^−1^, the maximum photosynthesis rate as well as the evapotranspiration rate of *G. avellana* was similar in low and high light acclimated plants; however, at higher PPFD the photosynthetic (A_max_ and WUE_i_) and photochemical (ΦPSII, ETR and qP) responses were significantly higher in high light acclimated plants, suggesting that *G. avellana* is able to grow under high light intensities without losing its intrinsic capacity to grow under low light. Our results also point out that this ability relies upon a flexible and dynamic molecular regulation of its photochemical and biochemical machineries, thus allowing plants to cope with the prevailing light environment in which they are growing.

Notable differences in the pattern of transcripts accumulation for a given light environment were revealed by the SOM clustering and allowed us to identify genes functionally relevant to photosynthesis and light-related process. SOM clustering has been used in many large-scale transcriptome data set analyses to detect groups of genes with similar expression profiles that yields biological relevant results (Chitwood et al., 2013, Ranjan et al., 2015, Ichihashi et al., 2014; Ostria-Gallardo et al., 2016). Because the use of block-wise module detection may influence the results and performance when there is a large number of blocks (van Dam et al., 2017), the use of SOM can help to obtain a reliable selection of a subset of genes in the transcriptome in order to construct a weighted gene co-expression network involved in light energy use for photosynthesis and related processes. The combination of WGCNA and the IGRAPH packages for network construction using this subset of genes produced a robust community structure whose properties and memberships were effective in identifying co-expression patterns with an appropriate biological relevance for the photochemical responses of *G. avellana*. In general, the use of gene co-expression networks has provided consistent relationships between transcriptional profiles and phenotypic traits (Zinkgraf et al., 2017; Suzuki et al., 2017). Our results revealed a dynamic molecular regulation of the photosynthetic machinery and carbon assimilation pathways as well as of the photoprotective and antioxidant system which is tightly coordinated with the prevailing light environment where this tree grows. Thus, differentially expressed genes included in module 1 belonged mostly to plants inhabiting low light environments and reflects co-expression of genes involved mainly in chlorophyll biosynthesis and regulation of the antenna complex size, reaction centers of photosystems, and oxygen evolving complex. By the other hand, genes included in modules four and fifteen belonged mostly to high light plants and were involved in carotenoid biosynthesis, respiration, cell wall formation, and photoprotection. Our *in-silico* construction of the light-driven gene co-expression network in *G. avellana*, when combined with an ecophysiological approach provides a comprehensive basis for understanding the mechanisms that allows this tree to have an optimal balance between photochemical, photoprotective and antioxidant performance. This occurs even in the extremes of the light gradient, as shown in our results for gas exchange and fluorescence of chlorophyll a, and ultimately, explains the acclimation allowing it to inhabit the wide range of light environments within the forest.

## Acknowledgments

E.O-G. thanks the Chilean National Commission for Scientific and Technological Research for doctoral fellowship, and the internship grant supported by Universidad de Concepción, project Mecesup UCO0708. Also thanks Dr. Carolina Sanhueza and Karina Rifo for their support in gas-exchange and fluorescence of chlorophyll *a* measurements, and Katalapi Park for excellent research field facilities. Part of the work was supported by NSF PGRP grant IOS—1238243 (to Julia Bailey-Serres, NRS, Siobhan Brady and Roger Deal).

## Author Contribution

E.O-G., L.J.C. and N.R.S. planned and designed the research. E.O-G. and L.J.C. conducted fieldwork and growth chamber experiments. E.O-G., A.R. and Y.I. performed bioinformatics, gene co-expression network, and data analysis. E.O-G. performed gas exchange and fluorescence of chlorophyll *a* analysis. E.O-G. and N.R.S. wrote the manuscript with contributions from other authors.

